# Epigenome-associated phenotypic acclimatization to ocean acidification in a reef-building coral

**DOI:** 10.1101/188227

**Authors:** Yi Jin Liew, Didier Zoccola, Yong Li, Eric Tambutté, Alexander A. Venn, Craig T. Michell, Guoxin Cui, Eva S. Deutekom, Jaap A. Kaandorp, Christian R. Voolstra, Sylvain Forêt, Denis Allemand, Sylvie Tambutté, Manuel Aranda

## Abstract

Over the last century, the anthropogenic production of CO_2_ has led to warmer (+0.74 °C) and more acidic (-0.1 pH) oceans^1^, resulting in increasingly frequent and severe mass bleaching events worldwide that precipitate global coral reef decline^2^,^3^. To mitigate this decline, proposals to augment the stress tolerance of corals through genetic and non-genetic means have been gaining traction^4^. Work on model systems has shown that environmentally induced alterations in DNA methylation can lead to phenotypic acclimatization^5,6^. While DNA methylation has been observed in corals^7-10^, its potential role in phenotypic plasticity has not yet been described. Here, we show that, similar to findings in mice^11^, DNA methylation significantly reduces spurious transcription in the Red Sea coral *Stylophora pistillata*, suggesting the evolutionary conservation of this essential mechanism in corals. Furthermore, we find that DNA methylation also reduces transcriptional noise by fine-tuning the expression of highly expressed genes. Analysis of DNA methylation patterns of corals subjected to long-term pH stress showed widespread changes in pathways regulating cell cycle and body size. Correspondingly, we found significant increases in cell and polyp sizes that resulted in more porous skeletons, supporting the maintenance of linear extension rates under conditions of reduced calcification. These findings suggest an epigenetic component in phenotypic acclimatization, providing corals with an additional mechanism to cope with climate change.

---

*Stylophora pistillata* is a globally distributed scleractinian coral with an available draft genome (Voolstra et al., 2017, under review). Previous work has demonstrated its plasticity and resilience in the face of high pCO_2_ conditions^12,13^. Remarkably, this coral remains capable of calcifying in seawater with significantly reduced pH of 7.2 even when aragonite, the main component of coral skeletons, falls below its saturation point in seawater (i.e., Ω_aragonite_ < 1)^13^. Previous work showed that the linear extension rate of *S. pistillata* in acidic seawater is not significantly different from that measured under control conditions (pH 8.0) despite the significantly reduced calcification rates^12^.

To investigate whether epigenetic mechanisms regulate phenotypic acclimatization to long-term pH stress, we cultivated *S. pistillata* colonies *in aquaria* for more than two years under four experimental conditions of seawater pH at 7.2, 7.4, 7.8 and 8.0 (control). Conditions in these aquaria were identical, except for the pCO_2_ and resulting differences in carbon chemistry in the tanks (3792, 2257, 856 and 538 μatm, respectively). We performed whole genome bisulphite sequencing on three replicate nubbins per tank and obtained data from 98% of all CpGs in the genome with a per-sample, per-position mean coverage of ~25 × (Supplementary Discussion 1, Supplementary Data 1).

The *S. pistillata* genome is sparsely methylated (1,406,097 bp, 7% of all CpGs), similar to other invertebrates, e.g., 1% in the bee *Apis mellifera*^14^; 2% in the wasp *Nasonia vitripennis*^15^; and 9% in the sea anemone *Nematostella vectensis*^16^. Like other invertebrates, the vast majority of cytosine methylation in *S. pistillata* occurs in genic (76.2%) rather than intergenic (22.9%) regions. Surprisingly, methylation in introns was higher than that in exons (Fig. 1a, 1b), unlike other invertebrates where introns have been reported to have low methylation^15,16^.

**Figure 1.**
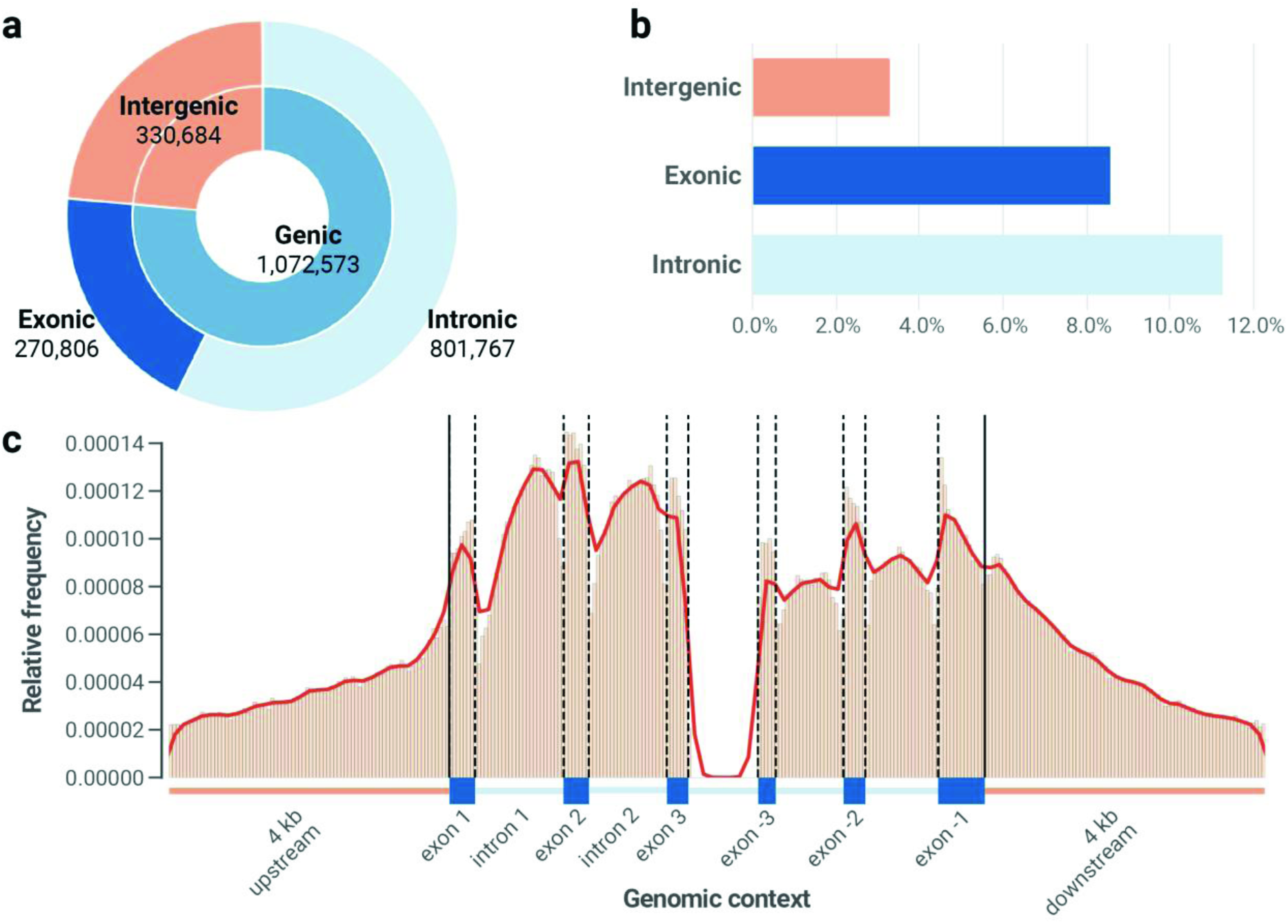
Epigenetic landscape of S. pistillata. (a) More than half of all methylated positions in *S. pistillata* are located in annotated introns. (b) Introns have proportionally more methylated positions (11.3%) than exons (8.6%) or intergenic regions (3.3%) even when accounting for the different amounts of CpG dinucleotides in the respective regions. (c) Relative frequencies of methylated positions across a standardised gene model with flanking 4 kb regions. Solid lines depict transcriptional start site (left) and transcription termination site (right). Exons and introns in the plot have normalised lengths that correspond to their respective mean lengths in *S. pistillata* (from left to right: 373 bp, 1,258 bp, 363 bp, 1,114 bp, 302 bp; 253 bp, 967 bp, 312 bp, 1,037 bp, 669 bp).

In contrast to vertebrates, methylation in the promoter regions of *S. pistillata* is scarce and does not seem to affect gene expression (see Supplementary Discussion 2); methylated positions are instead predominantly located within gene bodies (Figure 1c). Gene body methylation has recently been shown, in mice, to be established via crosstalk between the transcriptional machinery and histone modifications. RNA polymerase II-mediated transcription establishes new methylated positions along the gene body through the action of SetD2, H3K36Me3 and Dnmt3b. Methylation, in turn, reduces spurious transcription from cryptic promoters within these gene bodies^11^. Analyses of our RNA-seq data strongly indicate that these functions are conserved in corals. We find that gene body methylation increases with gene expression in *S. pistillata* (Fig. 2a, b), similar to other corals^7,8^. More importantly, we observe that methylated genes show significantly lower levels of spurious transcription relative to unmethylated genes (Fig. 2c). Furthermore, consistent with the repressive nature of methylation on expression, our analysis indicates that methylation in gene bodies also reduces transcriptional noise (lower variability of gene expression levels), echoing previous reports in corals and other organisms^7,17^ (Fig. 2d, Supplementary Discussion 3). These findings also suggest that the biological functions of gene body methylation are likely conserved across Metazoa, if at all present in the organism.

**Figure 2.**
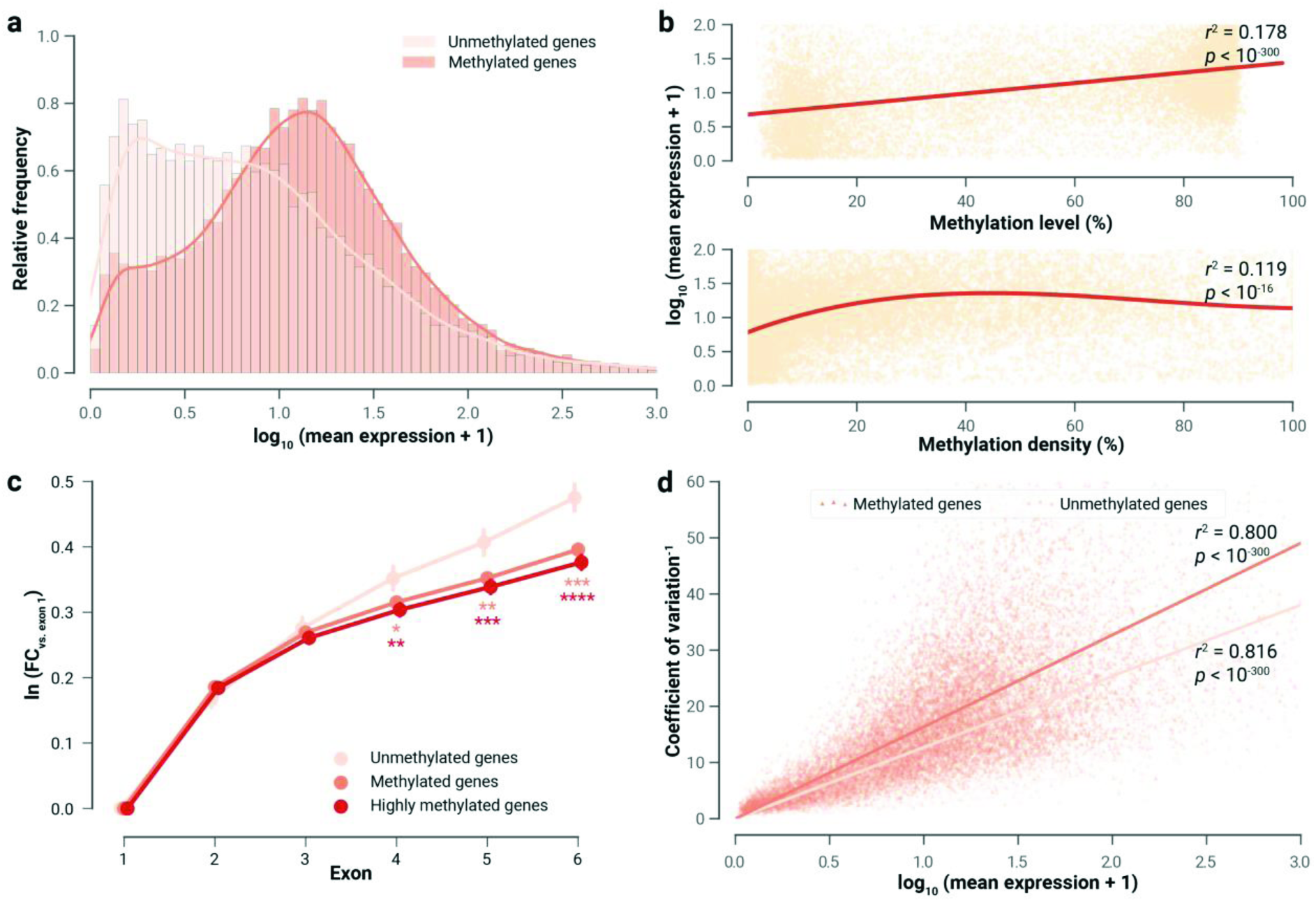
Effect of gene body methylation on genic expression. (a) The expression of methylated genes is significantly higher than that of unmethylated genes (*p* < 10^−300^, Student’s *t*-test). (b) Expression values for methylated genes are exponentially proportional to methylation level; however, the relationship of expression to methylation density is nonlinear. Methylation density at low levels is exponentially proportional to expression values, but it plateaus at ~40%. (c) Expression levels of the first six exons were calculated as natural logarithms of fold changes relative to the expression of the first exon. The difference in expression levels is likely driven by the reduction of cryptic transcription initiation in methylated genes. The difference is greater in highly methylated genes (median methylation level > 80%). Asterisks represent *p* values from *t*-tests of methylated (orange) or highly methylated genes (red) against unmethylated genes (peach), and coloured accordingly. *: *p* < 0.05; **: *p* < 0.01; ***: *p* < 0.001; ****: *p* < 0.0001. (d) There is a linear relationship between the inverse of the coefficient of variation (cv^−1^) and the log_10_-transformed mean expression values from all samples. The coefficient of variation, defined as the standard deviation of measured expression values divided by their mean, is consistently lower in methylated genes than unmethylated genes.

Based on these findings, we sought to further elucidate the potential role DNA methylation in phenotypic acclimation. Using generalised linear models (GLMs), we identified genes that undergo differential methylation in response to pH treatment (Supplementary Data 2). In general, we observed a significant increase in overall DNA methylation with decreasing pH, echoing similar observations in *Pocillopora damicornis*^9^. To validate these changes in methylation, we performed amplicon-specific bisulphite sequencing of selected genes on the original samples, as well as on samples of an independent experiment. These analyses showed high correlation of results obtained from whole genome and amplicon-specific bisulphite sequencing (*r*^2^ > 0.8, *p* < 0.01, Extended Data Fig. 5a) and further confirmed a high degree of reproducibility of DNA methylation changes across independent experiments (*p* < 0.01, Extended Data Fig. 5b). Analyses on laser-microdissected oral and aboral tissues further highlighted that most of the selected genes displayed strong and consistent tissue-specific methylation patterns, similar to findings in vertebrates^18^. These patterns were, in some cases, also correlated with known tissue-specific functions or activities of these genes (Supplementary Discussion 4).

Based on the skeletal phenotypes reported in previous studies analysing the effects of decreased pH levels on corals^12,19-21^, we initially expected that many of the differentially methylated and differentially expressed genes would be involved in biomineralization. However, among the calcification genes we characterized, relatively few genes responded. Overall, we observed a minor shift in methylation of organic matrix (OM) and extracellular matrix (ECM) components, with some known OM genes upmethylated and/or upregulated and others downmethylated and/or downregulated, implying a possible change in the scaffolding structure of the OM and ECM (Supplementary Discussion 5). As a general trend, we observed that Ca^2+^-binding OM constituents exhibited increased methylation and expression, whereas genes involved in cell-cell adhesion exhibited decreased methylation and expression.

Instead of biomineralization-related pathways, a functional enrichment analysis of differentially methylated genes revealed processes linked to growth and stress response (Fig. 3a, Supplementary Data 3). More specifically, we observed that the methylation levels of many genes in the MAPK signalling and cell growth pathways changed significantly (Fig. 3b). Remarkably, we found that the methylation levels of genes involved in the negative regulation of JNK and MAPK increased; conversely, the methylation levels of genes that positively regulate the same kinases decreased (Fig. 3c). For instance, the methylation of JIP1 (JNK-interacting protein 1, SpisGene10613), a negative regulator of JNK, increased in both aboral and oral tissues at pH 7.2; while the methylation of TRAF6 (TNF receptor-associated factor 6, SpisGene580), a positive regulator of JNK, decreased in both tissues. The JNK pathway has previously been shown to control cell, organ and body size in response to stress in mice^22^ and *Drosophila*^23^. The consistent differential methylation of JNK effectors that we observed in *S. pistillata* in response to decreasing pH levels therefore suggested a change in cell and body size.

**Figure 3.**
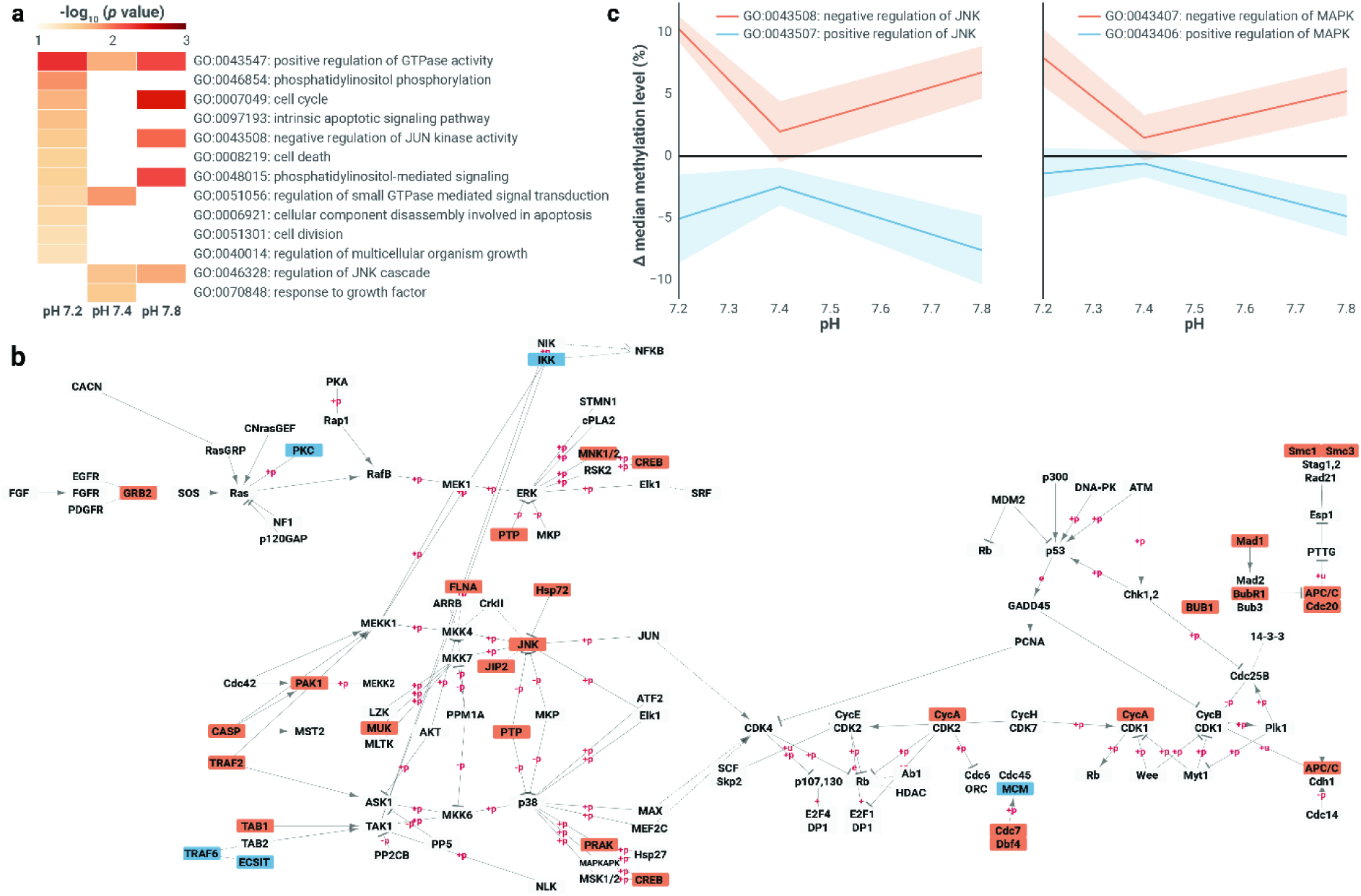
Differential methylation of genes in the MAPK/JNK signalling and cell growth pathways in the coral *S. pistillata* under pH stress. (a) The majority of the growth-and stress response-related GO terms are significantly enriched at pH 7.2. (b) At pH 7.2, more genes respond with a significant increase (red) than with a decrease (blue) in methylation; genes shaded in grey are methylated but not differentially regulated. (c) The methylation of genes that negatively regulate MAP kinase and JUN kinase (red lines) increases, whereas the methylation of genes that positively regulate the same kinases (blue lines) decreases in a reciprocal manner (error bars denote ±1 SE).

Analysis of differentially expressed genes in the MAPK and cell growth pathways highlighted several key genes that are differentially expressed in a manner that suggests a delay in cell division and an increase in cell size and body growth. For instance, Mos, a kinase that has been shown to prevent the degradation of cyclin B and thereby delay the onset of the anaphase^24^, was upregulated. Similarly, GADD45, a gene that binds to CDK1 and prevents its association with cyclin B, was also upregulated, potentially delaying the onset of M phase^25^. In contrast, Ras, which plays a key role in the progression of the G1 phase, was downregulated^26^. The kinases and phosphatases that regulate Ras experienced concomitant changes that indicated an overall reduction in Ras activity (Extended Data Fig. 1). For the latter, we observed expression increases in c-JUN, a transcription factor that promotes cellular growth; and Hsp70, previously observed in coral larvae under prolonged pH stress and presumably expressed to assist in the correct folding of cellular proteins^27^. The differential expression of these genes was subsequently corroborated via a quantitative reverse transcription PCR (RT-qPCR) analysis (Extended Data Fig. 2, Supplementary Data 4).

**Extended Data Figure 1.**
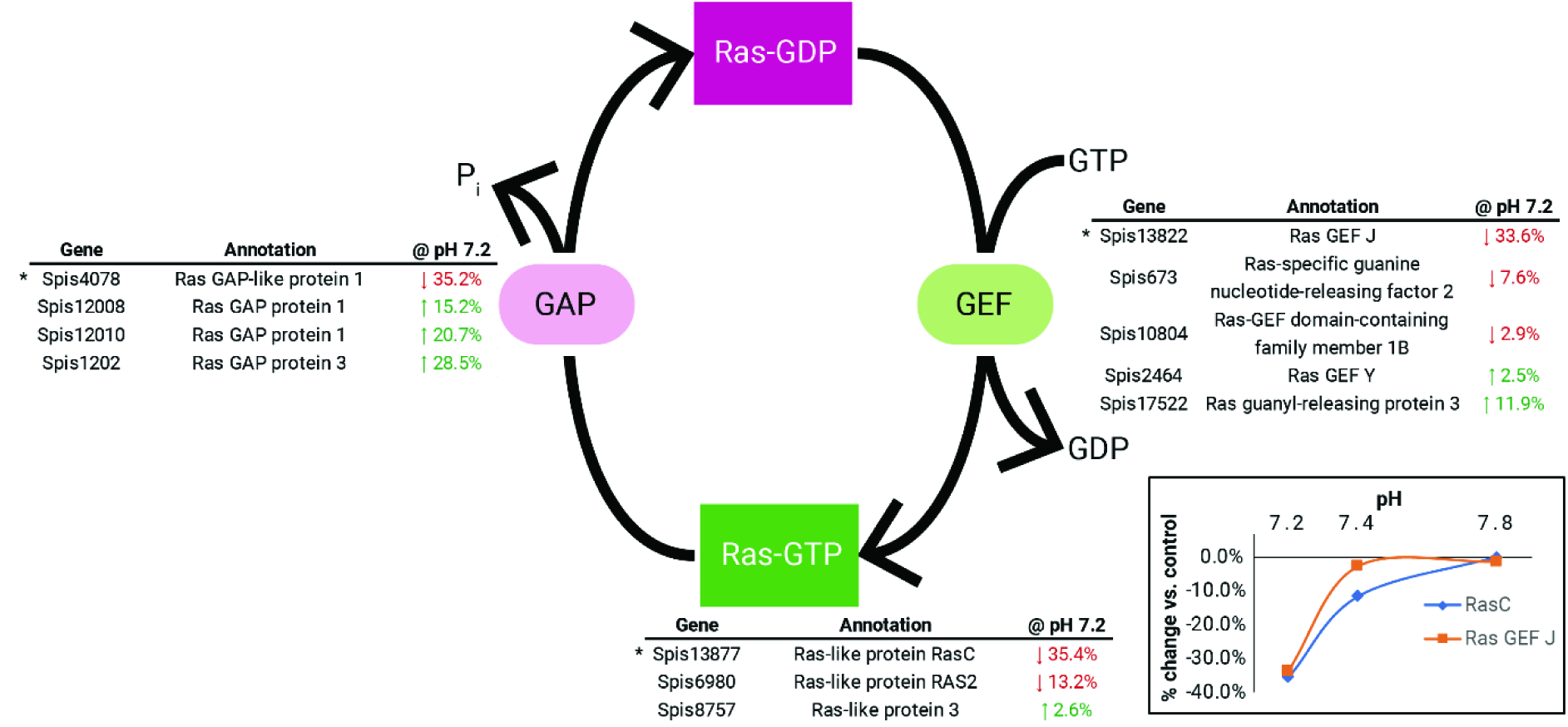
Downregulation of Ras and Ras guanine nucleotide-exchange factors (GEFs) and upregulation of Ras GTPase-activating proteins (GAPs) suggest a reduction in active Ras. The molecular switch regulating the activation of Ras is dependent on the activities of two opposing classes of proteins: GAPs, which convert the active Ras-GTP to the inactive Ras-GDP form, and GEFs which convert Ras-GDP to Ras-GTP. At pH 7.2, the Ras homologues present in *S. pistillata* (Spis8757, Spis6980 and Spis13877) had reduced expression. This occurred in tandem with the general upregulation of Ras GAPs and downregulation of Ras GEFs, thus further depleting the amount of active Ras-GTP under pH stress. Asterisks denote significant differential expression (*p* < 0.05). (inset) The significantly downregulated Ras homologue (Spis13877, blue) and Ras GEF (Spis13822, orange) exhibited differential expression exclusively at pH 7.2.

**Extended Data Figure 2.**
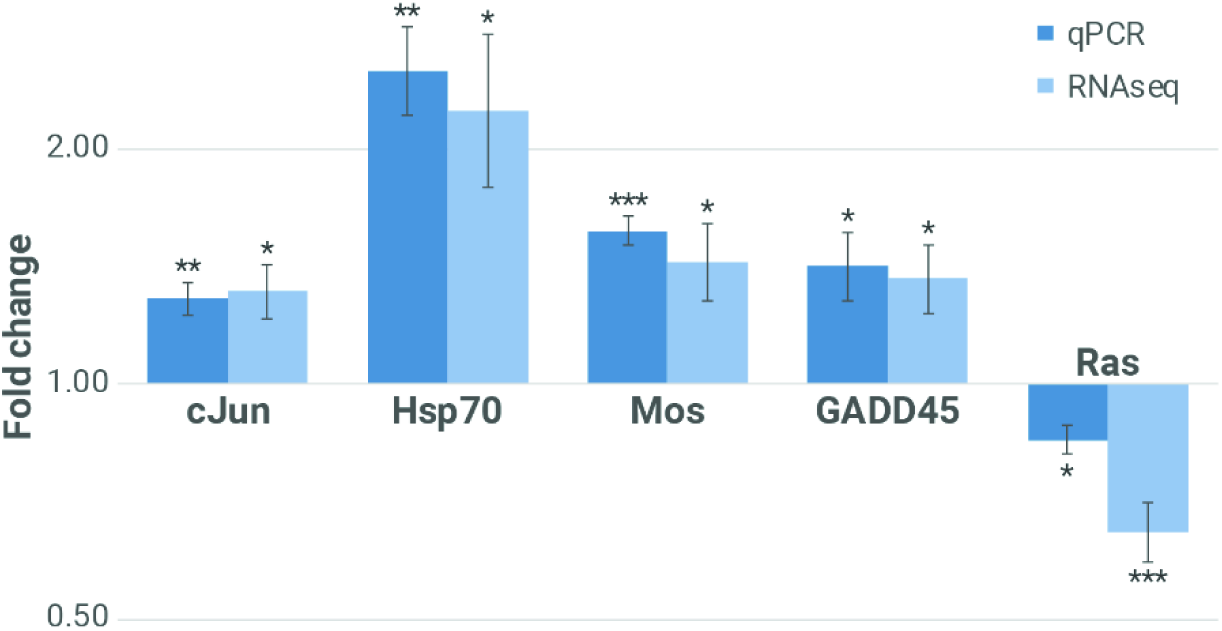
Differential expression of key genes was corroborated using RT-qPCR. Fold change indicates the expression of these genes at pH 7.2 relative to the control (pH 8.0). The experimentally measured values are very similar to that of RNA-seq, providing further support for the findings. Error bars represent ±1 SE. Asterisks denote significance of *t*-test *p* values. *: *p* < 0.05; **: *p* < 0.01; ***: *p* < 0.001.

In light of our data, we hypothesised that changes in DNA methylation (Fig. 4a) and expression of key genes lead to an increase in cell size at pH 7.2. We first confirmed that individual cell sizes were significantly larger in corals grown at pH 7.2 relative to the control (total *n* = 4,728 measurements across 14 nubbins, Fig. 4b, Supplementary Data 5a). We then hypothesized that this cell-level phenotypic change would be accompanied by a corresponding increase in polyp size, and ultimately, skeletal structure. Specifically, we posited that larger cells form larger polyps and larger corallite calyxes (the cup-shaped openings in the skeleton that house the polyps). As measuring the size of polyps is difficult due to their expandable/contractible nature, we opted to measure the size of the calyx as a proxy. As predicted, our analyses confirmed that the calyxes in corals grown at pH 7.2 relative to the control were indeed significantly larger (total *n* = 181 measurements across 10 nubbins, Fig. 4c, Supplementary Data 5b). Finally, we also confirmed that the skeleton was significantly more porous (*n* = 6 nubbins, Fig. 4d, e) in samples from the same treatment. All of the observed phenotypes were consistent with our hypothesis.

**Figure 4.**
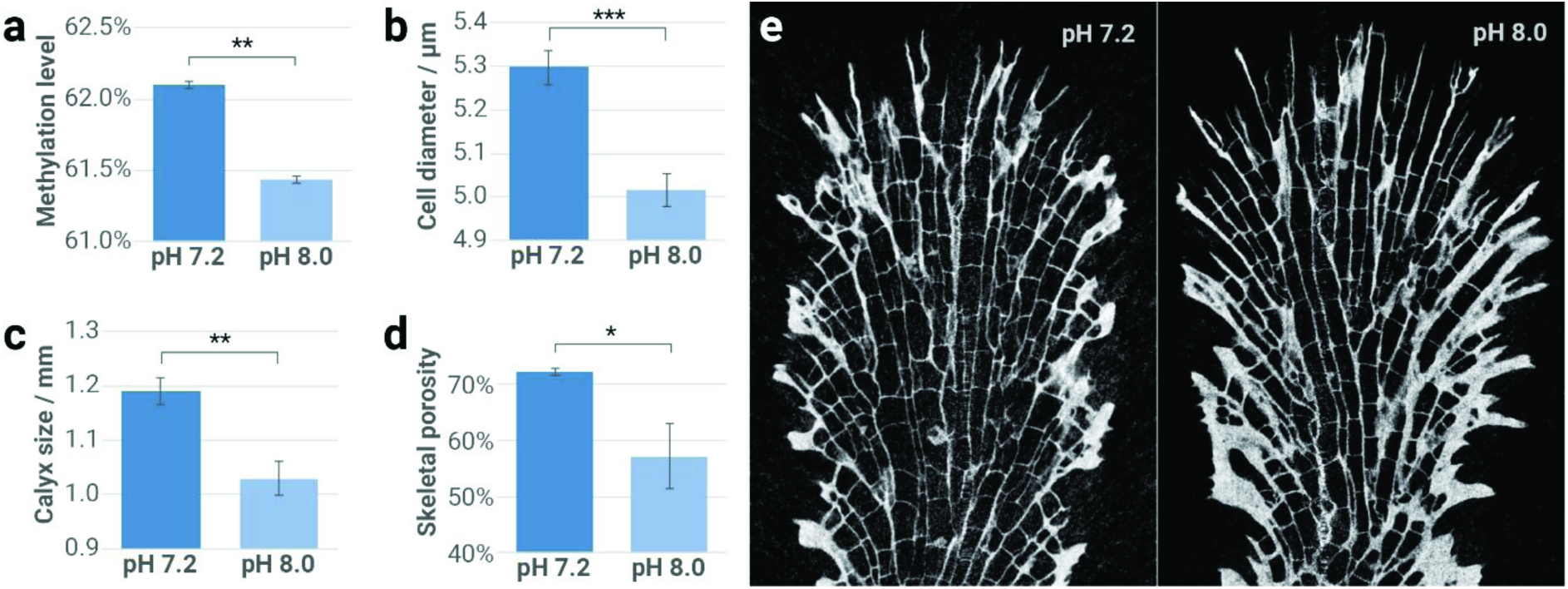
Effect of lower pH levels on cell growth. (a) Mean methylation levels were significantly increased at pH 7.2. (b) Cell sizes were significantly larger in nubbins grown in the pH 7.2 tank. (c) Larger cell sizes translated to larger polyps and consequently to a skeletal structure that contained larger calyxes. (d) The skeletal porosity was significantly higher at lower pH. (e) Representative longitudinal sections of *S. pistillata* skeletons under pH 7.2 and pH 8.0. Error bars represent 1 SE. Asterisks denote significance of *t*-test *p* values. *: *p* < 0.05; **: *p* < 0.01; ***: *p* < 0.001.

It is important to note that while our results show a strong correlation between changes in DNA methylation and the resulting phenotype, they do not show a *sensu stricto* causal relationship between changes in the methylation state and the phenotype. However, the high reproducibility of the changes in methylation and the presence of tissue-specific DNA methylation patterns lend support for a function of DNA methylation in phenotypic plasticity. Our results suggest that the observed phenotypic changes under pH stress are mediated through differential methylation and expression of known stress response pathways that control cell proliferation and growth^22,23^. We propose that these cellular phenotypic changes, together with shifts in organic matrix proteins, are among the drivers of morphological changes in the skeleton of this species under seawater acidification observed here and described previously^12^. These morphological changes towards a more porous skeleton are possibly a means by which *S. pistillata* can maintain linear extension rates in the face of depressed calcification rates under seawater acidification^12,28^. Such a trait would be advantageous in the benthic environment where competition for space and light is an important selective pressure^29^.

In conclusion, our results suggest that DNA methylation could offer corals greater ability to buffer the impacts of environmental changes and provide additional time for genetic adaptation to occur. Better understanding of the mechanisms underlying coral resilience will also provide additional avenues for reef-restoration efforts, such as the human-assisted acclimatization of corals in specialised nurseries (“designer reefs”^30^). Such efforts might prove crucial to averting large-scale losses of extant coral reefs in light of recent global declines due to climate change.

## Supplementary Discussion

### SD1. Comparisons with previous methylation studies in corals

In the absence of direct evidence, earlier work in corals^7,8^,^10^ based the identification of methylated genes on two assumptions. Firstly, methylated genes have coding sequences with fewer CpG dinucleotides than expected. This is usually quantified with CpG_O/E_ (also termed “CpG bias”), which represents the ratio of observed versus expected CpG dinucleotides on a per-gene basis. Secondly, strongly methylated genes have both high methylation levels (a particular CpG is methylated in most cells) and high methylation density (most CpGs in that gene are methylated).

Our data show that there are exceptions to both assumptions. A low CpG_O/E_ value (< 0.6) is fairly indicative of methylated genes, but classifying genes above the same threshold as unmethylated would include many genes that are methylated (Extended Data Fig. 3a). This could be from genes acquiring methylation at different evolutionary times and/or the presence of different selection pressures for methylated cytosines that undergo spontaneous deamination. Also, while correlations between CpG_O/E_, methylation level and methylation density in the expected direction exist, the strengths of the correlations are moderate at best (Extended Data Fig. 3b).

**Extended Data Figure 3.**
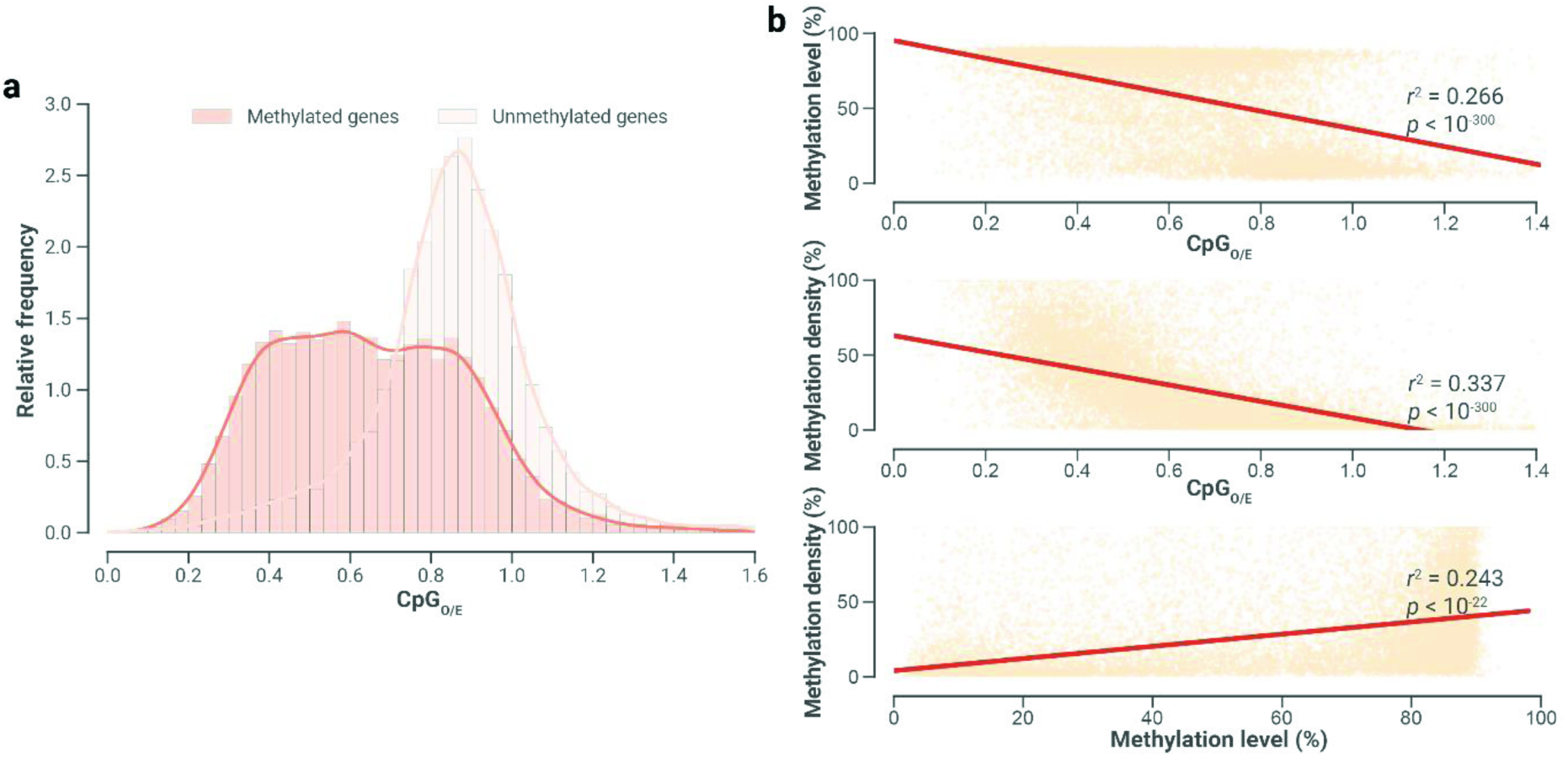
Distribution of CpG_O/E_ (“CpG bias”) for genes in S. pistillata. (a) While methylated genes do tend to have lower CpG_O/E_ values than non-methylated genes, a sizable fraction of methylated genes will be falsely considered non-methylated if a strict CpG_O/E_ threshold is used to define methylation states of genes. (b) Both metrics show moderate but statistically significant inverse correlation with CpG_O/E_. There is a moderately positive correlation between methylation level and methylation.

### SD2. S. pistillata promoter methylation has no effect on gene expression

Methylation in vertebrates has primarily been thought to drive gene expression via the differential methylation of promoter regions^31^. Our analysis indicates that–unlike in vertebrates–promoter methylation in *S. pistillata* is sparse and does not influence gene expression. We detected methylation in promoters (defined as a 4 kb window upstream of genes) of 6,675 genes (25.9% of all genes). We identified 720 expressed genes that had differential promoter methylation. Of these genes, only 20 genes were differentially expressed (*p* = 0.26, Fisher’s exact test), but only half of these genes exhibited the expected change in expression linked to the change in promoter methylation (Extended Data Fig. 4a). Also, methylated promoters are generally less methylated than are gene bodies, as evidenced through direct sequencing (Extended Data Fig. 4b) and CpG bias (Extended Data Fig. 4c), indicating that methylation was either established in gene bodies earlier in evolutionary time than in promoter regions, or gene bodies consistently have had higher levels of methylation than promoters have had.

**Extended Data Figure 4.**
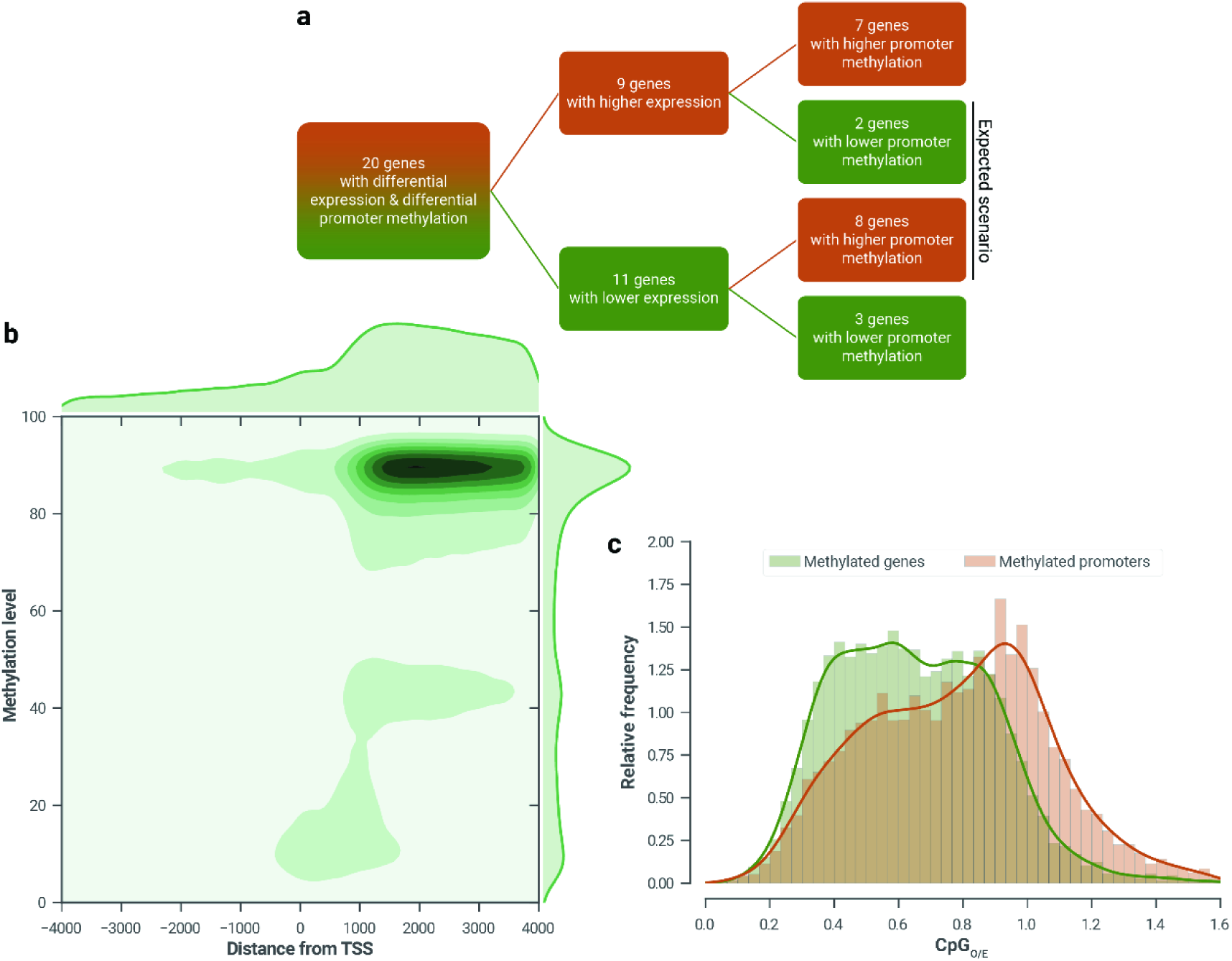
Multiple lines of evidence suggesting that promoter methylation does not influence expression patterns in S. pistillata. In these figures, promoter regions are defined as 4 kb windows upstream from the transcriptional start sites of all gene models. (a) Decision tree of 20 genes with differential gene expression and differential promoter methylation at pH 7.2 relative to the control. Only half of the genes exhibited the expected transcriptional response (i.e., increased promoter methylation represses expression and vice versa) (b) This heatmap of methylation levels in 4 kb windows upstream and downstream of transcription start sites demonstrates that methylation in promoter regions was much lower than that in gene bodies. (c) CpG_O/E_ values were much lower in gene bodies than in promoters.

### SD3. Correlation of genic expression to methylation

In light of the crosstalk between transcription, histone modifications and gene body methylation, in which highly expressed genes are methylated to suppress spurious transcription from cryptic promoters^11^, we sought to investigate whether similar observations apply to corals.

We first confirmed that expression of methylated genes is higher than unmethylated genes in *S. pistillata* (Fig. 2a). For methylated genes, expression levels were significantly higher in genes with higher methylation levels, and in genes with higher methylation density up to 40% (Fig. 2b). We also observed that genes that are highly methylated tended to be housekeeping genes (Supplementary Data 6b), in line with indirect evidence from three other corals from genus Acropora^7,8^.

To quantify the amount of spurious transcription, we calculated the expression of the first few exons relative to the first exon, with the expectation that methylation would reduce the expression of the middle exons. Our RNA-seq data showed that, for methylated genes, there was a progressive reduction of internal expression across the genes, culminating in a significant difference observed in exon 6. The decline was also more significant when we restricted the analysis to highly methylated genes (Fig. 2c).

Some reports have suggested a link between increased methylation and the reduction in transcriptional variability (i.e., transcriptional noise)^7,17^, estimated as the coefficient of variation among the measured gene expression values. As previous studies did not investigate whether expression values could be a confounding factor for transcriptional noise, we sought to model transcriptional noise as a function of both expression level and methylation state. Our GLM analysis indicates that while expression level largely (73.4-74.3%, *p* < 0.0001) determines transcriptional noise, the presence of methylation in a gene further reduces (25.7-26.6%, *p* < 0.001) the noise, consistent with a suppressive action of methylation on expression (Extended Data Table 1).

**Extended Data Table 1.** Modelling transcriptional noise as a function of expression level and methylation state. In our GLM analysis, the inverse of the coefficient of variation (cv^−1^) was correlated against methylation status (0 for unmethylated genes; 1 for methylated genes) and log_10_-transformed expression values. Among the tested variables, expression had the greatest determinant of transcriptional noise (11.74), followed by an expression-dependent coefficient that is present only in methylated genes (4.44). Methylation state, on its own, has close to negligible effect on transcriptional noise (-0.19). All variables significantly influence transcriptional noise (*p* < 0.001).

|  |  |  |  |  | 95% confidence interval |  |
| --- | --- | --- | --- | --- | --- | --- |
|  | Coefficient | Standard error | z | P>\|z\| | [0.025 | 0.975] |
| Intercept | 1.006 | 0.0349 | 28.8416 | 0.0000 | 0.9376 | 1.0743 |
| expression | 11.7377 | 0.089 | 131.8157 | 0.0000 | 11.5632 | 11.9122 |
| methylation_state | -0.1942 | 0.0553 | -3.5079 | 0.0005 | -0.3026 | -0.0857 |
| expression:methylation_state | 4.4417 | 0.1253 | 35.4583 | 0.0000 | 4.1962 | 4.6872 |

### SD4. Validation of methylation patterns

To validate our initial findings and their reproducibility, we used MiSeq-based amplicon-specific bisulphite sequencing of selected candidate genes on the original samples as well as on samples from an independent repeat experiment.

Accuracy of amplicon-specific bisulphite sequencing was assessed by performing this technique on DNA from all three biological replicates from the pH 7.2 and control treatments that were used to construct the original whole genome bisulphite libraries. Methylation levels of individual CpGs assayed using both techniques were highly correlated *(r^2^ >* 0.8, *p <* 0.01, Extended Data Fig. 5a, Supplementary Data 7). For most of the designed amplicons, both techniques produced concordant predictions of the shift in methylation state at pH 7.2 (binomial test *p*(X ≥ 14) = 0.002, Extended Data Fig. 5b).

**Extended Data Figure 5.**
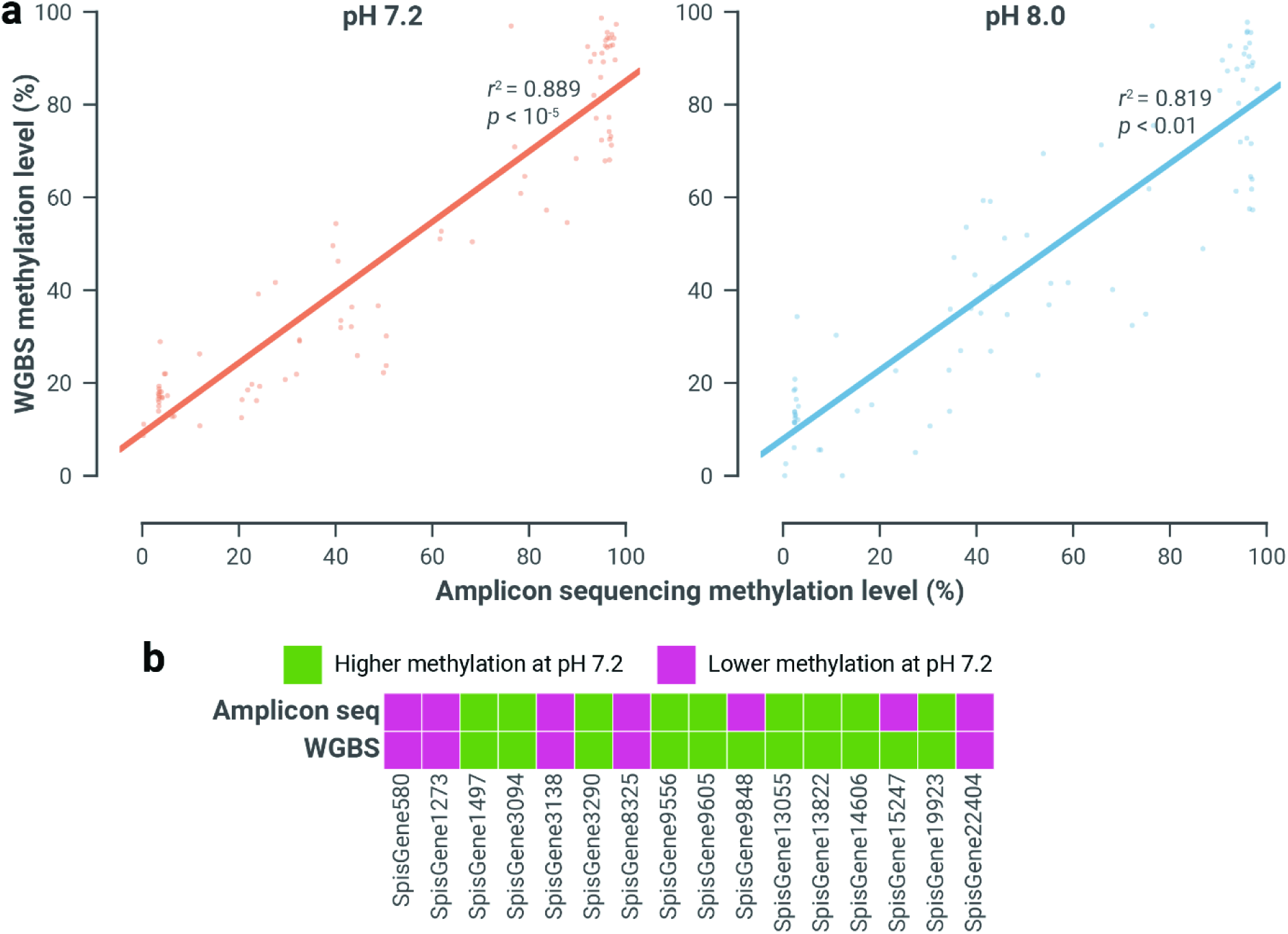
Amplicon-specific bisulphite sequencing can accurately assay methylation levels of amplicons of interest. (a) Results from amplicon sequencing largely corroborated that from whole genome bisulphite sequencing (*r*^2^ > 0.8 and *n* = 3 per treatment). (b) It also produced the expected methylation pattern in most of the tested amplicons (14 of 16 genes, binomial test *p*(X ≥ 14) = 0.002).

To verify that the observed methylation changes are also reproducible, and at the same time investigate whether methylation changes exhibited tissue-specific patterns, we performed laser microdissections on samples from an independent experiment with the same conditions as the previous pH 7.2 and 8.0 samples. We separated the aboral tissues from the oral tissues (Extended Data Fig. 6a), and subsequently carried out amplicon-specific bisulphite sequencing on the extracted DNA. Of the 20 genes tested, amplicon sequencing confirms that 15 genes were differentially methylated in the expected direction (Extended Data Fig. 6b, Supplementary Data 8, binomial test *p*(X ≥ 15) = 0.02).

**Extended Data Figure 6.**
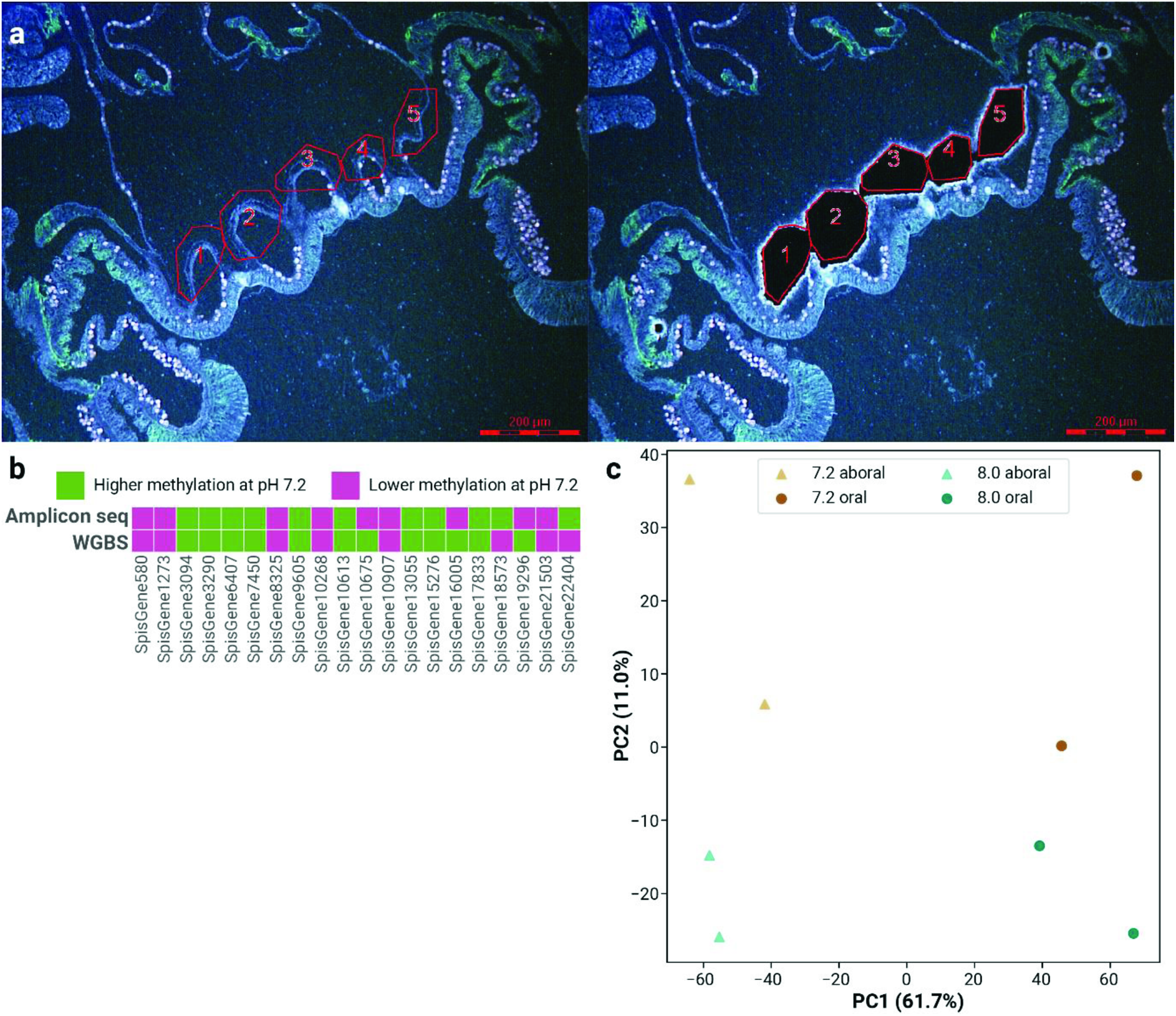
Methylation patterns are strongly tissue-specific. (a) Example laser microdissection of aboral tissue from a fixed sample (scale bar represents 200 µm). (b) 15 out of 20 genes (binomial test *p*(X ≥ 15) = 0.02) showed the expected methylation patterns (i.e., loci that had higher/lower methylation at pH 7.2) in samples from a separate experiment. (c) PCA on these loci show that methylation patterns have a very strong tissue-specific signature (along PC1); less so by treatment (along PC2).

Interestingly, we find that tissue-specific methylation patterns are stronger than treatment-specific patterns (Extended Data Fig. 6c). This observation ties in well with studies done on humans^18^, where non-cancerous cell lines tended to cluster based on tissue of origin. Nonetheless, the effect of the long-term pH stress is evident: the second principal component of the Principal Component Analysis (PCA) cleanly separates the treatments from each other.

Among the tested amplicons, some had similar methylation levels in both tissues, while stark differences were present in some others. For example, major yolk protein (Spis7450), which is found in the skeletal organic matrix, has a mean methylation level of 26.2% in aboral tissues, in contrast to the 8.5% in oral tissues. Catalase (SpisGene3094), which has previously been associated with symbiosis, has a 13.6% methylation in aboral tissues, a third of the 43.3% in the oral tissues where most of the symbionts reside.

Unexpectedly, genes that are involved in the regulation of JNK also displayed tissue-specificity. TRAF6 (SpisGene580) and JNK-interacting protein 1 (SpisGene10613) were far more methylated in oral tissues than aboral. This potentially indicates that these genes might be regulated in a tissue-specific manner.

### SD5. Differential expression and methylation of biomineralization genes

pH stress has been consistently linked to the differential regulation of biomineralization genes in scleractinian corals^32-34^; however, its effect on methylation patterns has yet to be studied. Our analysis of differentially methylated and differentially expressed biomineralization genes did not correlate well with the observed phenotype. Also, the corresponding methylation landscape of these genes was strikingly different from that of expression (Extended Data Fig. 7, Supplementary Data 9). This indicates that pathways responding to pH stress are perhaps under direct expression regulation via transcriptional factors, rather than affected by the fine-tuning of expression afforded via changes in methylation state.

**Extended Data Figure 7.**
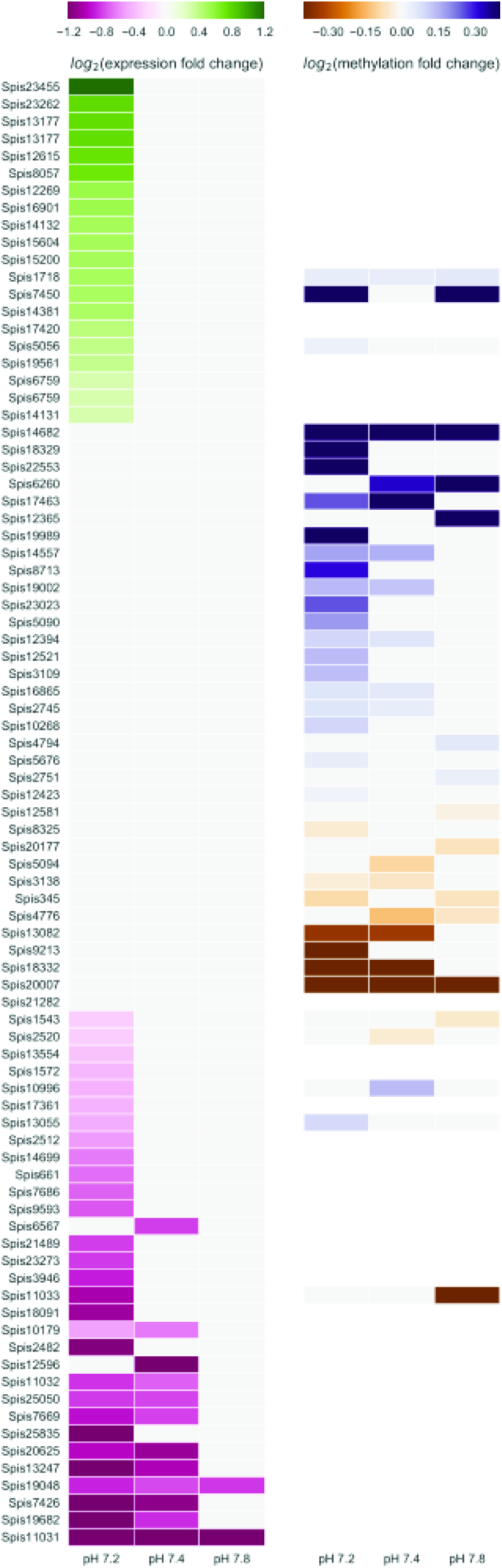
Effect of long-term pH stress on selected biomineralization-related genes. All genes are significantly differentially expressed and/or methylated (*p* < 0.05) in one or more experimental conditions. Genes are sorted by the mean increase in expression, followed by the mean increase in methylation. Grey boxes represent changes in expression or methylation that were not statistically significant; blank boxes represent unmethylated genes. The heatmaps show little overlap between differentially expressed genes and differentially methylated ones, but the expression and methylation responses were always more pronounced at lower pH.

Among the differentially methylated genes, we identified two genes encoding key ion transporters putatively involved in the calcification process. The first is a gene encoding the carbonic anhydrase STPCA (Spis16865) that has previously been localized in the ECM and is thought to facilitate calcification by hydrating local CO_2_ to HCO_3_^-^ ^32^. The gene for this protein was more methylated in the pH 7.2 and 7.4 treatments, indicating a potential compensating mechanism that increases HCO_3_^-^ concentration. The second gene encodes the bicarbonate transporter SLC4ß (Spis8325), which is thought to be coupled to the enzyme STPCA2 in calicoblastic cells to facilitate HCO_3_^-^ transport to the ECM^35^. This gene, however, is less methylated at pH 7.2, potentially indicating a modulation of bicarbonate ion transport to the ECM.

Interestingly, we also observed increased expression of CARP1 (Spis6759), a main constituent of the OM. This gene has been shown to be important for mineral deposition^36^ and is localized in the cells around the skeleton^37^. CARPs and analogous proteins can bind to collagen—the former serving as mineral nucleation points, while the latter providing structural support within the ECM. An increase in CARP transcription may indicate an increase in possible nucleation points for growth of the coral skeleton and might represent a compensation mechanism. Major yolk protein (MYP, Spis7450), another OM protein with elevated expression^36^, is unusual in also having increased methylation at pH 7.2. MYP binds and shuttles ferric iron^38^, which is an important trace element in the coral skeleton and potentially plays a photoprotective role^39^. Furthermore, we found a putative homologue of bone morphogenetic protein (BMP1, Spis8057) that was upregulated at pH 7.2. This protein was also found to be highly expressed in embryos of the sea urchin *Strongylocentrotus purpuratus* immediately before primitive skeleton (spicule) formation^40^, possibly indicating a positive role in calcification.

## Methods

### Growth conditions of S. pistillata

Colonies of the tropical coral *Stylophora pistillata* were exposed to long-term seawater acidification as described previously ^12,13^. Briefly, corals were kept *in aquaria* supplied with Mediterranean seawater (exchange rate of 70% h^−1^) at a salinity of 38 g L^−1^, temperature of 25 °C and irradiance of 170 mmol photons m^−2^ s^−1^ on a 12h/12h photoperiod provided by HQI-10000K metal halide lamps (BLV Nepturion, Steinhoering, Germany). Carbonate chemistry was manipulated by bubbling CO_2_ to reduce the control pH (pH 8.0) to the target values of pH 7.8, 7.4, and 7.2. Values of carbonate chemistry parameters were as previously measured: 3792, 2257, 856 and 538 μatm respectively for pH 7.2, 7.4, 7.8 and 8.0^12^.

### Identification of methylated CpGs

DNA was extracted from *S. pistillata* nubbins (triplicates of four growth conditions) using a nuclei isolation approach to minimize contamination with symbiont DNA, as previously described (Voolstra et al., 2017, in review). Briefly, cells from *S. pistillata* from a nubbin of about 3 cm were harvested using a Water Pick in 50 ml of 0.2 M EDTA solution refrigerated at 4 °C. Extracts were successively passed through a 100 µm and a 40 µm cell strainer

(Falcon, Corning, NY) to eliminate most of the algal symbionts. Extracts were then centrifuged at 2,000*g* for 10 min at 4 °C. The supernatant was discarded and the resulting pellets were homogenized in lysis buffer (G2) of the Qiagen Genomic DNA Isolation Kit (Qiagen, Hilden, Germany). The DNA was extracted following manufacturer instructions using Genomic-tip 100/G (Qiagen, Hilden, Germany). DNA concentration was determined by O.D. with Epoch Microplate Spectrophotometer (BioTek, Winooski, VT). Contamination with *Symbiodinium* DNA was assessed via PCR targeting the multicopy gene RuBisCO (Genbank accession number AY996050).

Bisulphite DNA libraries were prepared following a modified version of the NEBNext Ultra II DNA Library Prep Kit for Illumina (NEB, Ipswich, MA). Methylated TruSeq Illumina adapters (Illumina, San Diego, CA) were used during the adapter ligation step followed by bisulphite conversion with the EpiTect Bisulfite kit (Qiagen, Hilden, Germany), with the following cycling conditions (95 °C for 5 min, 60 °C for 25 min, 95 °C for 5 min, 60 °C for 85 min, 95 °C for 5 min, 60 °C for 175 min, then 3 cycles of 95 °C for 5 min and 60 °C for 180 min. Hold at 20 °C ≤ 5 hours). The final library was enriched with the KAPA HiFi HotStart Uracil+ ReadyMix (2X) (KAPA Biosystems, Wilmington, MA) following the standard protocol for bisulfite-converted NGS library amplification. Final libraries were quality checked using the Bioanalyzer DNA 1K chip (Agilent, Santa Clara, CA), and quantified using Qubit 2.0 (Thermo Fisher Scientific, Waltham, MA), then pooled in equimolar ratios and sequenced on the HiSeq2000 platform. Sequencing of the libraries resulted in 1.53 billion read pairs across 12 samples (Supplementary Data 1). The raw sequences were trimmed using cutadapt v1.8^41^. Trimmed reads were then mapped to the *S. pistillata* genome, deduplicated and scored on a per-position basis for methylated and unmethylated reads using Bismark v0. 13^42^.

To ensure that methylated positions were *bona fide*, three stringent filters were used. Firstly, the probability of methylated positions arising from chance on a per-position basis was modelled using a binomial distribution B(*n*, *p*), where *n* represents the coverage (methylated + unmethylated reads) and *p* the probability of sequencing error (set to 0.01 to mimic a Phred score of 20). We kept positions with *k* methylated reads if *p*(X ≥ *k*) < 0.05 (post-FDR correction). Secondly, methylated positions had to have at least a methylated read in all three biological replicates of at least one growth condition. Finally, the median coverage of the position across all 12 samples had to be ≥ 10. These steps ensured that methylated positions were highly replicable and highly covered.

### Assignment of genomic context to methylated cytosines

Based on the genome annotation of *S. pistillata* (in the form of a GFF3 file) and the positional coordinates of the methylated cytosines (in a tab-separated file produced by Bismark), a Python script was written to annotate every methylated cytosine with additional genomic context. The script determined whether the methylated position resides in a genic or intergenic region: for the former, an additional check occurs to determine whether it is in an exon or an intron. Subsequently, distances to the 5’ and 3’ end of each genomic feature (gene/intergenic region/exon/intron) were calculated.

### Identification of differentially methylated genes

Using the methylation level of genes at pH 8.0 as a control, GLMs were implemented in R to identify genes that were differently methylated at pH 7.2, 7.4 and 7.8 respectively. The general formula used was:

> glm(methylated, non_methylated ~ pH * position, family=“binomial”)

where “methylated, non_methylated” was a two-column response variable denoting the number of methylated and non-methylated reads at a particular position, while “pH” and “position” were predictor variables for the pH of the environment and the genomic coordinate of the position respectively. Data from individual replicates were entered separately—this approach assigned equal weightage to each replicate, instead of having a disproportionate skew towards the replicate with the highest coverage if the data were pooled. Genes with < 5 methylated positions were filtered out to reduce type I errors.

### Identification of differentially methylated promoters

As promoter regions were not defined in the *S. pistillata* genomes, it was assumed to be located in a window of 4 kb upstream of the transcription start site. A GLM similar to the one described in the previous paragraph was used to identify differentially methylated promoters; however, due to the scarcity of methylated positions in these windows, genes with ≥ 2 methylated positions were retained for this analysis.

### Identification of differentially expressed genes

High quality total RNA was extracted for library creation from 3 *S. pistillata* nubbins per treatment. Directional mRNA libraries were produced using the NEBNext Ultra Directional RNA Library Prep Kit for Illumina (NEB) as described in (Voolstra et al., 2017, in review). A total of 674 million paired-end reads (read length of 101 bp) were retrieved from 6 lanes on the HiSeq2000 platform (Illumina, San Diego, CA).

Trimming was intentionally left out to increase the number of mapped reads, and to reduce bias^43^. The resulting 362 million trimmed reads were mapped to *S. pistillata* gene models with kallisto v0.42.4^44^ to produce TPM (transcripts per million reads) values. Based on these values, sleuth^45^ was used to identify differentially expressed genes by contrasting all biological replicates of pH 7.2, 7.4 and 7.8 against the controls (pH 8.0).

### Calculation of exon expression relative to first exon

As kallisto is a pseudo-mapper, i.e., it assigns reads to gene models, but not the exact location of where the read maps to the gene model, reads from all 12 replicates were mapped separately using HISAT2 (v2.1.0)^46^ against the *S. pistillata* genome. Genomic positions that correspond to exonic locations were extracted with a Perl script, and mean coverages were computed to produce RPKM values on a per-replicate, per-exon basis with a Python script.

Genes with six or more exons were selected for the analysis. Furthermore, as ratio-based computations are skewed by lowly expressed genes, genes with overall mean RPKM > 0.5 and non-zero expression in the first six exons across all 12 replicates were selected. Ratios were subsequently natural log-transformed to improve normality, resulting in 1,141 unmethylated genes, 5.750 methylated genes, and 2,955 highly methylated genes (genes with median methylation > 80%).

### Functional enrichment of methylated genes

GO term annotations were obtained from literature (Voolstra et al., 2017, in review). GO term enrichment analyses were carried out with topGO^47^ with default settings. GO terms with *p* < 0.05 and occurring ≥ 5 times in the background set were considered significant. Multiple testing correction was not applied on the resulting *p*-values as the tests were considered to be non-independent^47^.

KEGG Orthology (KO) annotations were merged from results of gene model annotation and KAAS (KEGG Automatic Annotation Server), http://www.genome.jp/tools/kaas/, with parameters “GHOSTZ”, “eukaryotes”, and “bi-directional best hit”. Based on the KO annotation, KEGG pathway enrichment was carried out using Fisher’s exact test. Pathways with *p* < 0.05 were considered significant.

### RT-qPCR verification of key growth genes

There is only one reference amplicon (against β-actin) designed specifically for this organism^32^. As normalization against a single reference gene is rarely acceptable^48^, additional control genes were selected from our RNA-seq data. Selected genes were highly abundant (TPMs > 1,000), and expressed at roughly equivalent levels in all sequenced libraries.

Primers were designed, whenever possible, to include a large intronic region in addition to an exonic size of ~100 bp. A Python script that interfaced with Primer3 v2.3.6^49^ was written to optimise primers design, e.g., melting temperatures of ~60 °C, GC% of 30-70%, avoid long runs of a single base and no strong secondary structure.

A preliminary RT-PCR was run to confirm that amplicons produced band sizes corresponding to the amplified exonic region; as expected, none of the primers produced detectable bands that included the intronic regions. Six reverse transcription reactions (on total RNA from all pH 7.2 and 8.0 replicates) were carried out with SuperScript III First-Strand Synthesis SuperMix (Invitrogen, Carlsbad, CA) with the supplied oligo-dT primers. The subsequent qPCR was carried out using Platinum SYBR Green qPCR SuperMix-UDG (Invitrogen, Carlsbad, CA) in a 7900HT Fast Real-Time PCR System (Applied Biosystems, Waltham, MA). Both protocols were carried out according to manufacturer’s instruction.

Primer sequences, amplicon sizes, and the analysis of the RT-qPCR results are fully detailed in Supplementary Data 4.

### Laser microdissection of *S. pistillata* samples

Apexes of colonies were prepared as described previously^50^. Briefly, apexes of *S. pistillata* from an independent experiment using the same treatment conditions were fixed in 3% paraformaldehyde in S22 buffer (450 mM NaCl, 10 mM KCl, 58 mM MgCl_2_, 10 mM CaCl_2_, 100 mM Hepes, pH 7.8) at 4 °C overnight and then decalcified, using 0.5 M ethylenediaminetetraacetic acid (EDTA) in Ca^2+^-free S22 at 4 °C. They were then dehydrated in an ethanol series and embedded in Paraplast. Cross-sections (6 µm thick) were cut and mounted on POL-membrane (0.9 µm) frame slides (Leica Microsystems, Wetzlar, Germany). The Leica AS LMD system, with a pulsed 337 nm ultraviolet laser on an upright microscope, was used for the microdissections. The laser beam can be moved with a software-controlled mirror system that allows selecting target cells and tissues. Target cells can be preselected on a monitor with a freehand drawing tool, and then the computer-controlled mirror moves the laser beam along the pre-selected path and the target cells are excised from the section. The dissected part then falls into a PCR tube under gravity. The collection by gravity ensures quick and contamination-free processing of the dissected tissue sections.

DNA from eight sections (duplicates of aboral and oral tissue from pH 7.2 and from controls) were extracted, and subjected to bisulphite conversion using EpiTect Plus Bisulfite Conversion kit following manufacturer’s instructions (Qiagen, Hilden, Germany).

### Validation of differential methylation via amplicon-specific bisulphite sequencing

As dissected tissues contain minute amounts of DNA, we decided to perform amplicon-specific bisulphite sequencing to validate the methylation levels within our amplicons of interest. A nested PCR design was used to generate these amplicons. Outer primers were pooled in the initial PCR run (35 cycles), and the resulting reaction mix was then evenly split into amplicon-specific individual PCR reactions (35 cycles) with their respective inner primers. As the amplicons were to be sequenced on the Illumina MiSeq platform, the inner primer pairs were designed with additional overhangs to facilitate downstream library creation, per the 16S Metagenomic Sequencing Library Preparation guide (Illumina, San Diego, CA).

Amplicons were selected within genes that exhibited differential methylation at pH 7.2 relative to control. As bisulphite-conversion produces Watson and Crick strands that are no longer reverse complements of each other, primers were designed to amplify the sense strand of the gene product. To optimise primer design, a self-written Python script that interfaced with Primer3 v2.3.6^49^ was used to select primer pairs that were fully located within non-methylated regions. This important consideration avoids the need for degenerate primers (one degenerate base is required per methylated CpG), reducing ordering costs and dodging potential PCR amplification biases. In total, 46 amplicons (i.e., 184 primers) were designed.

To assess the primers, test nested PCRs were performed on converted *S. pistillata* total DNA; PCR products were subsequently visualized on an agarose gel. Primers that completely failed to produce amplicons with expected band sizes were discarded (*n* = 3). The remaining primers were grouped into 3 batches (of ~14 primers each) to reduce unintended products that might arise from pooling too many outer primers in the initial reaction.

To validate our original findings and to test the accuracy of this approach, the same amplicon-specific bisulphite approach was carried out on DNA from all three samples of pH 7.2 and control that were used to construct the original whole genome bisulphite libraries. This allowed for direct comparisons of methylation levels assayed via amplicon-specific and whole genome bisulphite sequencing.

With the use of Nextera XT indices (Illumina, San Diego, CA), libraries were pooled and sequenced on one-and-a-half MiSeq runs. A total of 14.2 million paired-end reads (read length of 300 bp) were produced. These reads were cleaned and mapped to the *S. pistillata* genome with the same pipeline used to process reads from whole genome bisulphite sequencing, with the sole exception of skipping the deduplication step (distinct amplicons from the same loci map to the same genomic coordinates and thus are erroneously deemed duplicates).

We used a very conservative coverage threshold for the downstream analyses: only methylated positions with read coverages greater than 100 in all samples were retained. This filtering step increased the precision of the measured methylation levels and reduced the effect of noise on the results.

### Measurement of cell sizes

As described previously^28^, branches of 2-3 cm size were sampled from colonies grown in pH 7.2 and 8.0 treatment and placed in a 7% MgCl_2_ solution to anesthetize tissues. Oral discs (the apparent portion of the polyp) were then cut from the colony under a microscope using microdissection scissors with 5 mm blades. Cells were then dissociated using a sterile tube pestle and homogenized by repeated pipetting using a 200 µl pipette. Suspended cells (50 µl) were mounted between the slide and coverslip, and 30 random pictures of the surface were taken. Cell sizes on these pictures were subsequently measured using SAISAM software (Microvision Instruments, France). In all, 4,728 measurements were taken across 14 tested nubbins (Supplementary Data 5a).

### Measurement of corallite calyx size

Branches of similar size (2 cm in length) were sampled from colonies in the pH 7.2 and 8.0 treatments and placed in a 10% NaClO solution for 2 h to remove tissues. Skeletons were rinsed several times in tap water, followed by ultra-pure H_2_O, and then dried at 40 °C for at least 24 h. Samples (5 replicates per pH condition) were coated with gold/palladium and observed at 4 kV with a JEOL 6010LV electron microscope. Diameters of the corallite calyxes were measured using manufacturer-provided SMile View software (JEOL, Tokyo, Japan). A total of 181 measurements were performed across 10 branches (Supplementary Data 5b).

### Measurement of skeletal porosity

The non-invasive, high-resolution imaging method of X-ray micro-computed tomography (micro-CT) was used as described to measure skeletal porosity^12^. Micro-CT analysis was carried out at the Polyclinique St Jean, Cagnes sur Mer, France, with an SkyScan 1173 compact micro-CT (SkyScan, Antwerp, Belgium). A microfocus X-ray tube with a focal spot of 10 mm was used as a source (80 kV, 100 µA). The sample was rotated 360° between the X-ray source and the camera. Rotation steps of 1.5° were used. At each angle, an X-ray exposure was recorded on the distortion-free flat-panel sensor (resolution 2,240 × 2,240 pixels). The resulting slice was made of voxels, the three-dimensional equivalent of pixels. Each voxel was assigned a grey value derived from a linear attenuation coefficient that relates to the density of materials being scanned. All specimens were scanned at the same voxel size. The radial projections were reconstructed into a three-dimensional matrix of isotropic voxels ranging from 5 to 10 mm, depending on the exact height of the coral tip. X-ray images were transformed by NRecon software (SkyScan) to reconstruct two-dimensional images for quantitative analysis. From these images, evaluation of the morphometric parameters was performed using CT-Analyser software (SkyScan). A manual greyscale threshold was implemented manually on the first set of images and then applied to all specimens. For each sample, a digital region of interest was created to extend through 100 µm of skeleton at 7 mm distance from the apex, corresponding to about 15 slices. The percentage of negative space in relation to the skeleton was then determined, providing the measure of porosity. For each treatment condition, three branches of similar size were taken from the apical part of a colony. Porosity was analysed in one experimental trial with three replicates per treatment.

## Accession codes

### Whole genome bisulphite sequencing and transcriptomic data

PRJNA386774.

## Acknowledgements

We thank Dominique Desgre, Natacha Caminiti-Segonds and Nathalie Techer for assistance in coral husbandry; the KAUST Sequencing Core Facility for the sequencing of the libraries; Nathalie Techer for cell size measurements; and Pierre Alemanno and Christophe Sattonnet (Polyclinique St Jean, Cagnes sur Mer, France) for access to the micro-CT. This publication is based upon work supported by the King Abdullah University of Science and Technology (KAUST) Office of Sponsored Research (OSR) under Award No. FCC/1/1973-22-01.

## Author information

**Affiliations**

**Red Sea Research Center, King Abdullah University of Science and Technology, Saudi Arabia**

Yi Jin Liew, Yong Li, Craig T. Michell, Guoxin Cui, Christian R. Voolstra & Manuel Aranda

**Centre Scientifique de Monaco, Marine Biology Department, Principality of Monaco**

Didier Zoccola, Eric Tambutté, Sylvie Tambutté, Denis Allemand & Alexander A. Venn

**Computational Science Lab, Faculty of Science, University of Amsterdam, Amsterdam, The Netherlands**

Eva S. Deutekom & Jaap A. Kaandorp

**Passed away 17 December 2016**

Sylvain Forêt

## Contributions

M.A. conceived and coordinated the project. M.A., C.R.V., D.Z., E.T., S.T., D.A. and A.A.V provided tools, reagents and/or data. C.T.M. constructed libraries for whole genome bisulphite sequencing and RNA-seq. Y.J.L., Y.L., E.S.D., J.A.K. and M.A. analysed expression data. Y.J.L, S.F. and M.A. analysed methylation data. Y.J.L., D.Z., G.C. and M.A. performed tissue-specific experiments. E.T. and A.A.V. performed and analysed skeleton parameters measurements. Y.J.L. and M.A. wrote the manuscript. All authors except for S.F. (passed away) read and approved the final manuscript.

## Corresponding author

Correspondence to: Manuel Aranda.

